# Serological evidence for localized and persistent antibody response in Zika virus-positive neonates with microcephaly (Brazil, 2015)- a secondary analysis

**DOI:** 10.1101/051326

**Authors:** Andrew A. Lover

## Abstract

A recent publication in *The Lancet* by Cordeiro and colleagues reported levels of IgM for Zika (ZIKV) and dengue (DENV) viruses in serum and cerebrospinal fluid (CSF) of 31 infants born with microcephaly in Brazil.^1^ Their study suggests higher titers in CSF relative to serum in individual neonates, but no quantitative comparisons were reported. In this short report, the differences in antibody titers are quantified and compared between sample sources; across sampling periods; and between sample sources within individual neonates to more comprehensively understand these data to inform serological surveillance. These are statistically significant differences in ZIKV titers between CSF and serum samples, (in contrast to DENV titers), and these ZIKV titer levels remain elevated across sampling dates, whereas the titer in serum trends downward by sampling date. In multivariate models, ZIKV titer in CSF samples is *independent* of titer in serum, and of DENV antibodies in both CSF and serum. These findings quantify the compartmentalization of ZIKV antigens across the blood-brain barrier, and suggest complex interplay between ZIKV and cross-reacting DENV antigens in congenital/neonatal infections.

**Note:** Dataset for analysis appears to Appendix II.

## I. Introduction

Zika virus (ZIKV) has recently emerged from relative obscurity to the forefront of the global health agenda in 2015 due to a potential association with microcephaly and other congenital malformations.^2^

Diagnosis of ZIKV can be difficult due to a short window to capture active viremia, and cross-reactivity with other flaviviruses including Yellow fever virus (YFV) and dengue (DENV) in many serological assays. Several sets of recent case reports have specifically highlight the difficulties with both IgG and IgM cross-reactivity using ELISA.^3,4^

Several studies have examined the serology of ZIKV in adults, including pregnant women, but there has been limited data from neonates. The data reported by Cordeiro *et.al* fill an important gap; and cursory examination of their data suggest that there are differences in measured viral titers between sample sources from individual neonates, and between the cross-reacting antigens from a pool of all four DENV serotypes. In this report, statistical comparisons are made to more comprehensively understand these data and the potential relationships between measured serological parameters for these flaviviruses in the context of microcephaly in Brazil.

## II. Methods

Full details of the patient population, and IgM serological assays are available in the original publication by Cordeiro *et al*.; data were extracted from the table.^1^

Comparison of population-level optical density (OD) ratios (titers) for each paired sample type was via a non-parametric signrank test. For multivariate models, the outcomes of ZIKV and DENV titers in CSF were used due to the inherent challenges in obtaining these samples relative to serum samples.

For reported regression coefficients, univariate models residuals were examined for outliers using Cooks’ distance and leverage. All included variables were ‘forced’ into the model without explicit model building, but were examined for collinearity. Due to several outliers, reported R^2^ values are from robust regressions;^5^ however all conclusions are unchanged from ordinary least-squares (OLS) regressions. For exploratory trends lines, lowess (locally weighted scatterplot smoothing^6^), a non-parametric trend line method used with a bandwidth of 0.9.

All analyses used Stata 14.1 (College Station, TX, USA). All tests were two-tailed, with α = 0.05; no corrections were made for multiple comparisons.

## III. Results

Comparisons of the population-level optical densities as median values for ZIKV and DENV (<u>Figure 1</u>, and <u>Table 1</u>) show no evidence of a statistically significant difference between DENV titer in the CSF and serum samples (p = 0.345). However comparison of the ZIKV antibodies in the same population shows evidence for a statistically significant difference (p = 0.016) between the two sets of paired CSF and serum samples. These differences are also clearly evident in <u>Figure 1</u>, with little variance in the optical densities ratios for DENV, but higher values and greater variability in measured ZIKV titers.

**Figure 1.**
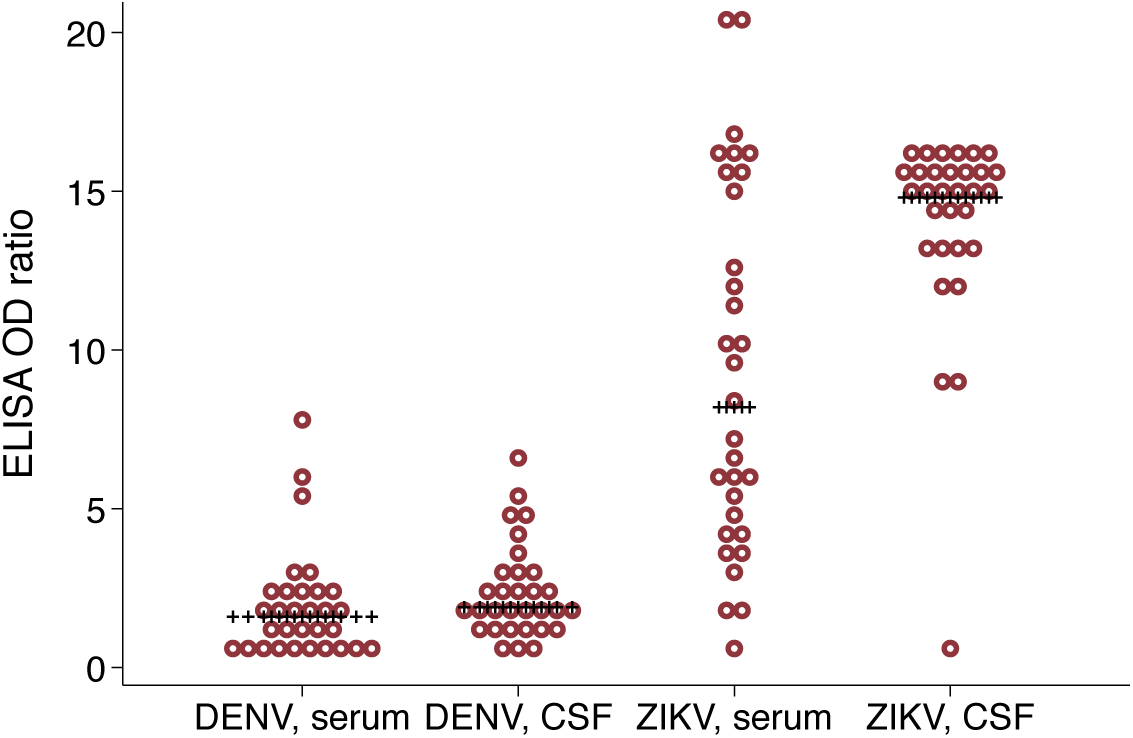
<u>Distribution of titers for DENV and ZIKV in neonates by sample source, Brazil 2015. Note: median values marked; (N= 31).</u>

**Table 1.** <u>Comparison of median measured optical density ratios by sample source, Brazil 2015.</u>

| Comparison | Test | Full data<br>(N=31)<br>p-value<br>for difference | ZIKV(+) only<br>(N=30)<br>p-value<br>for difference |
| --- | --- | --- | --- |
| ZIKV in CSF vs.<br>ZIKV in serum | signrank | 0.029 | 0.016 |
| DENV in CSF vs.<br>DENV in serum | signrank | 0.265 | 0.345 |

Correlations between the titer of each sample type by antigen are show in <u>Figure 2</u>, with best-fit linear regression lines. There is a clear positive relationship between DENV titers in serum and CSF (R^2^ = 0.31, but there is no clear relationship between ZIKV titers in serum and CSF (R^2^ = 0.002).

**Figure 2.**
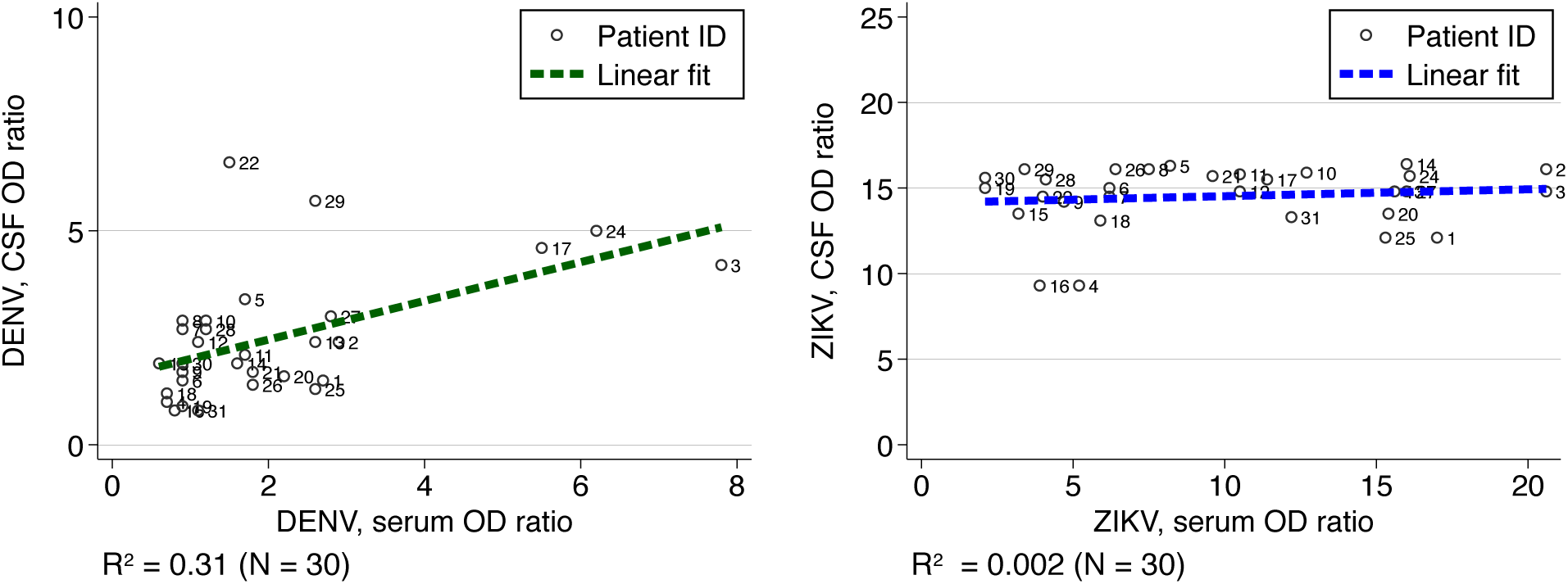
<u>Correlations between sample sources in measured serological titers for DENV and ZIKV in neonates, Brazil 2015. Note: R^2^ from robust regressions.</u>

Plots of titers by sampling date with non-parametric trend lines suggest comparable levels of DENV IgM in CSF and serum samples (<u>Figure 3</u>, <u>left panel</u>). Conversely, comparison of the ZIKV IgM levels suggests divergent trends (<u>Figure 3</u>, <u>right panel</u>). While the ZIKV IgM in serum trends downward by sampling date, the titer in CSF remains consistently high in all neonates independent of the sampling date. A complementary set of comparisons of titers between DENV and ZIKV within each sampling type (<u>Appendix figure 1</u>) suggests proportional levels of both IgMs within the CSF and serum samples across sampling time.

**Figure 3.**
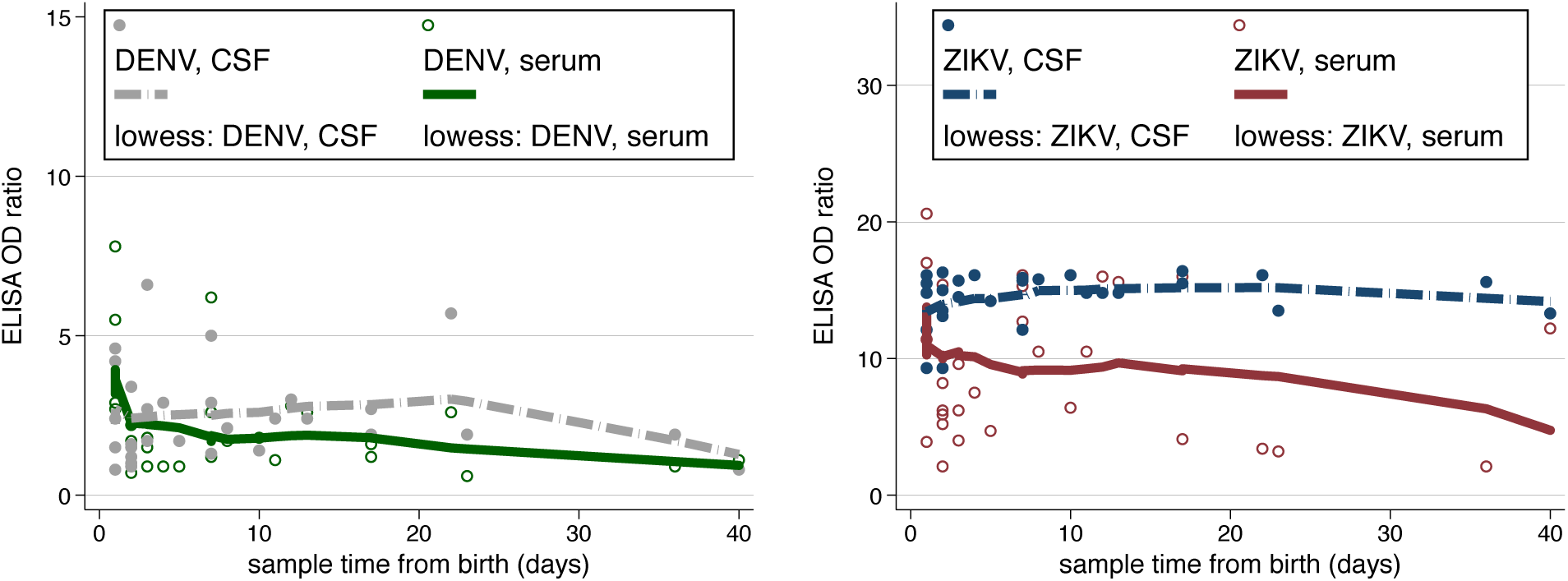
<u>Comparison of DENV and ZIKV IgM levels in neonates by sample date, with local smoothing trend lines (lowess), Brazil 2015 (N= 30) Note change of scale between panels.</u>

Univariate and multivariate analysis of ZIKV and DENV titers in CSF are presented in <u>Tables 3 and 4</u>. With adjustment for sampling day after birth, the measured DENV titers in CSF have a positive statistically significant relationship with the ZIKV in CSF and DENV in serum, and a negative statistically significant relationship with ZIKV in serum.

**Table 3.** <u>Univariate and multivariate predictors of measured **ZIKV** antibody OD in CSF samples (N= 30). Notes: factors in boldface are significant at p < 0.05. Multivariate model R^2^ = 0.14.</u>

| Univariate |  |  | Multivariate |  |  |
| --- | --- | --- | --- | --- | --- |
| Factor | Coefficient<br>(95% CI) | p-value | Factor | Coefficient<br>(95% CI) | p-value |
| **DENV in CSF** | 0.428<br>(0.042 to 0.815) | 0.030 | DENV in CSF | 0.540<br>(-0.616 to 1.70) | 0.360 |
| ZIKV in serum | 0.012<br>(-0.078 to 0.102) | 0.795 | ZIKV in serum | 0.037<br>(-0.109 to 0.183) | 0.615 |
| DENV in serum | 0.138<br>(-0.502 to 0.325) | 0.151 | DENV in serum | -0.153<br>(-0.855 to 0.550) | 0.670 |
| Days from birth | 0.007<br>(-0.0434 to 0.057) | 0.789 | Days from birth | 0.018<br>(-0.033 to 0.69) | 0.492 |
|  |  |  | constant | 13.153<br>(10.361 to 15.944) | > 0.001 |

After adjustment for covariates, this model predicts that for each single unit increase in DENV in CSF optical density ratio there will be a 0.277 (95% CI: 0.166 to 0.388, p < 0.001) unit increase in ZIKV in CSF; a 0.082 (95% CI: - 0.153 to -0.011, p = 0.024) unit decrease in ZIKV in serum; and a 0.558 unit (95% CI: 0.376 to 0.741, P < 0.001) increase of DENV in serum. Days from birth was not significantly associated with DENV titer in CSF, with a coefficient of -0.007 (95% CI: -0.030 to 0.016, p = 0.553).

In sharp contrast to these results, for the outcome of ZIKV titer in CSF, none of the exploratory factors were significantly associated with outcome in multivariate models (all p > 0.30).

## IV. Conclusions

This secondary analysis examines the relationship between ELISA titers for both DENV and ZIKV in paired serum and CSF samples. These results suggest contrasting compartmentalization of antibodies across the blood brain barrier, with statistically significantly higher levels of ZIKV antibodies in the CSF relative to serum in individual neonates.

Moreover, a limited exploratory analysis suggests these levels may remain higher in the weeks post-birth, assuming that each subject’s levels are comparable at birth, while cross-reacting DENV titers in the CSF appear to decay through time.

The most striking finding is the independence of ZIKV titers in the CSF, whereas cross-reacting DENV antigens are closely correlated with other antigens in both sample types. This may be due to a ceiling effect in the serological assays, which could limit the dynamic range of these data in the absence of further serial dilutions. However, even with this caveat in mind, the titer levels in CSF are ca. 5- to 10-fold higher than those in serum, and ca. 15-fold higher than DENV titers in the CSF. This finding suggests that ZIKV titer in the CSF cannot be estimated directly from the titer in serum, or from cross-reacting DENV titers using this CDC assay at these dilutions.

There are limitations in this analysis: the sample size is limited; the subject inclusion/exclusion criteria are uncertain; other pathogens that may cause congenital malformations may not have been tested for; and a large number of potential covariates are unavailable (especially maternal IgM or IgG; maternal age; and gestational age).

Complex dynamics between serological assays for flaviviruses have been reported at the population level in other settings e.g.^7^ and greater understanding and quantification of these trends may inform both ZIKV neuropathology studies and serological surveillance in areas with multiple co-circulating arboviruses.

---

**Table 2.** <u>Univariate and multivariate predictors of measured **DENV** antibody OD in CSF samples (N= 30). Notes: factors in boldface are significant at p < 0.05. Multivariate model R^2^ = 0.49.</u>

| Univariate |  |  | Multivariate |  |  |
| --- | --- | --- | --- | --- | --- |
| Factor | Coefficient<br>(95% CI) | p-value | Factor | Coefficient<br>(95% CI) | p-value |
| **ZIKV in CSF** | 0.303<br>(0.178 to 0.428) | < 0.001 | **ZIKV in CSF** | 0.277<br>(0.166 to 0.388) | < 0.001 |
| ZIKV in serum | 0.047<br>(-0.025 to 0.120) | 0.201 | **ZIKV in serum** | -0.082<br>(- 0.153 to -0.011) | 0.024 |
| **DENV in serum** | 0.461<br>(0.299 to 0.622) | < 0.001 | **DENV in serum** | 0.558<br>(0.376 to 0.741) | < 0.001 |
| Days from birth | -0.011<br>(-0.049 to 0.027) | 0.568 | Days from birth | -0.007<br>(-0.030 to 0.016) | 0.553 |
|  |  |  | constant | -1.920<br>(-3.033 to -0.806) | 0.001 |

**Appendix figure 1.**
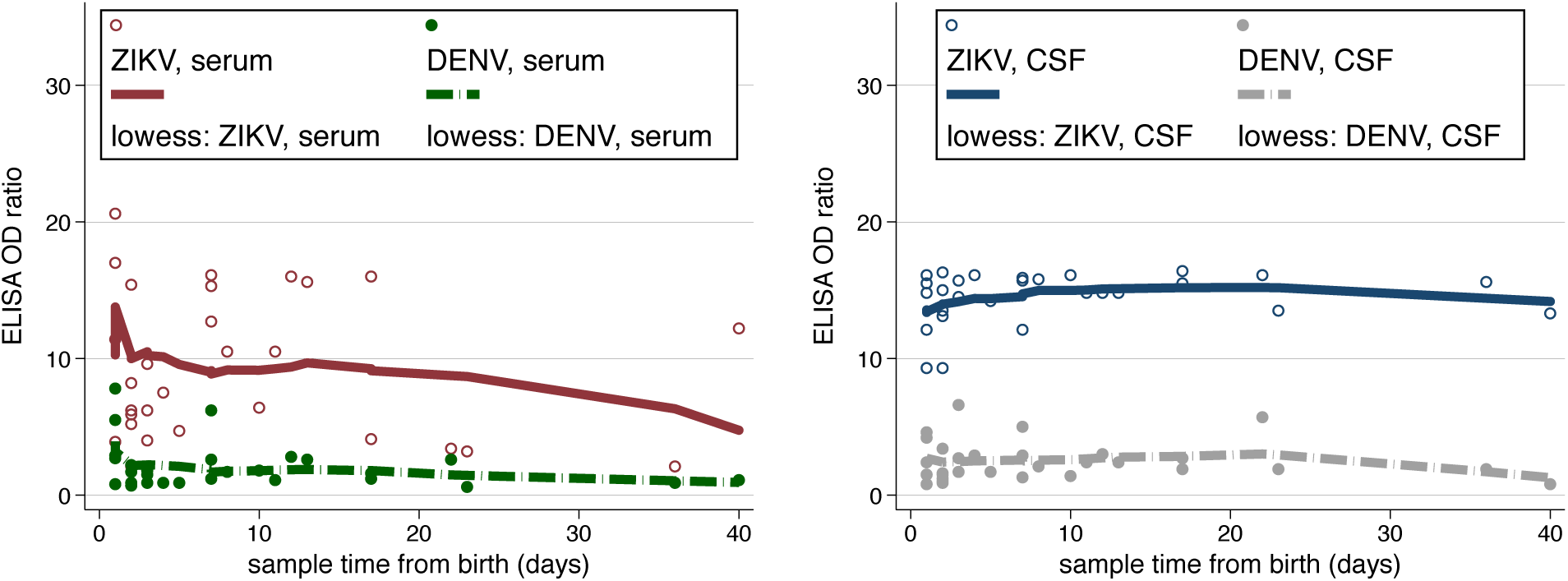
<u>Comparison of DENV and ZIKV IgM levels in neonates by sample date, across sample location with local smoothing trend lines (lowess), Brazil 2015 (N= 30).</u>

**Appendix figure 2.**
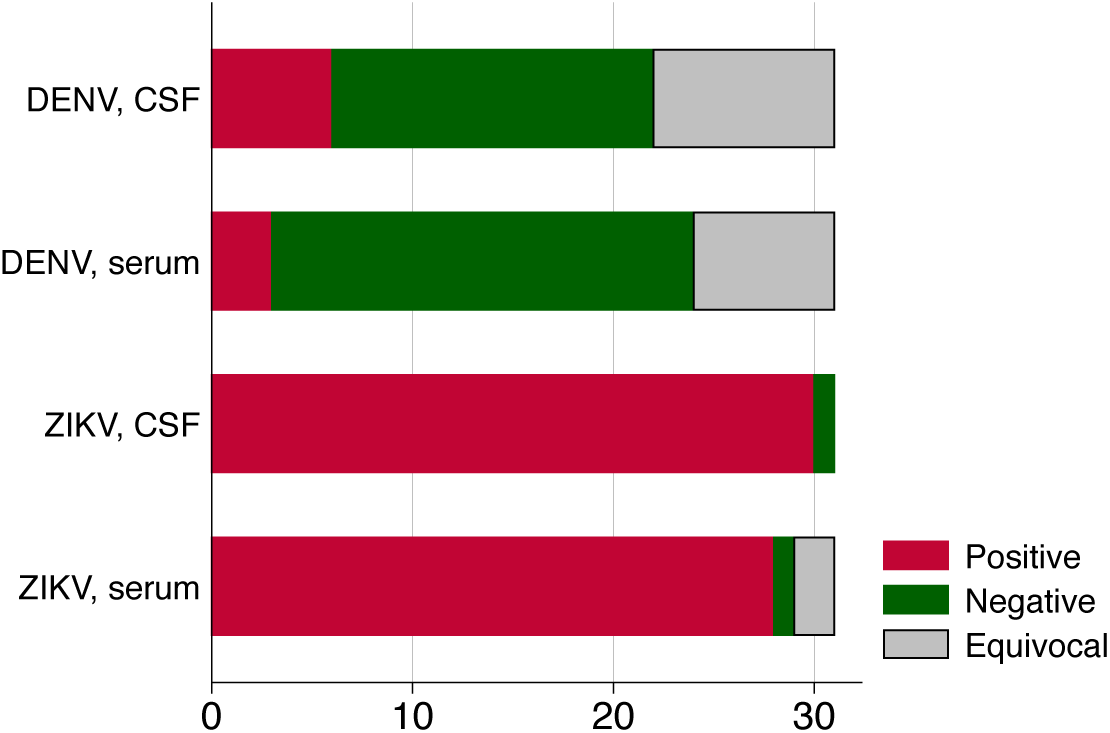
<u>Seropositive, seronegative, and equivocal proportions in neonatal serology, Brazil 2015 (N= 30) Note: cutoffs as in reference^1^</u>

**Appendix II.**
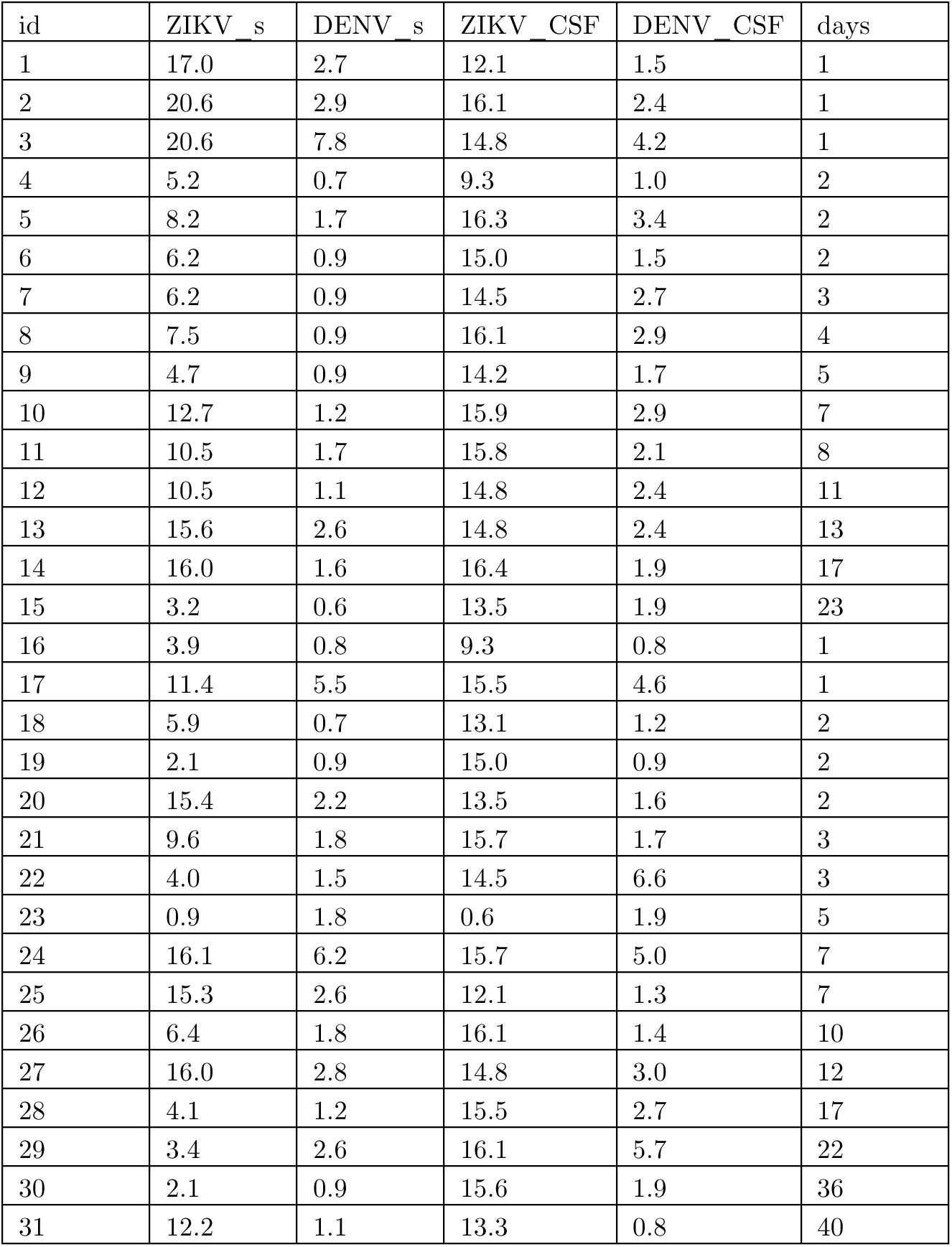
<u>dataset for analysis</u>

| id | ZIKV _s | DENV _s | ZIKV _CSF | DENV _CSF | days |
| --- | --- | --- | --- | --- | --- |
| 1 | 17.0 | 2.7 | 12.1 | 1.5 | 1 |
| 2 | 20.6 | 2.9 | 16.1 | 2.4 | 1 |
| 3 | 20.6 | 7.8 | 14.8 | 4.2 | 1 |
| 4 | 5.2 | 0.7 | 9.3 | 1.0 | 2 |
| 5 | 8.2 | 1.7 | 16.3 | 3.4 | 2 |
| 6 | 6.2 | 0.9 | 15.0 | 1.5 | 2 |
| 7 | 6.2 | 0.9 | 14.5 | 2.7 | 3 |
| 8 | 7.5 | 0.9 | 16.1 | 2.9 | 4 |
| 9 | 4.7 | 0.9 | 14.2 | 1.7 | 5 |
| 10 | 12.7 | 1.2 | 15.9 | 2.9 | 7 |
| 11 | 10.5 | 1.7 | 15.8 | 2.1 | 8 |
| 12 | 10.5 | 1.1 | 14.8 | 2.4 | 11 |
| 13 | 15.6 | 2.6 | 14.8 | 2.4 | 13 |
| 14 | 16.0 | 1.6 | 16.4 | 1.9 | 17 |
| 15 | 3.2 | 0.6 | 13.5 | 1.9 | 23 |
| 16 | 3.9 | 0.8 | 9.3 | 0.8 | 1 |
| 17 | 11.4 | 5.5 | 15.5 | 4.6 | 1 |
| 18 | 5.9 | 0.7 | 13.1 | 1.2 | 2 |
| 19 | 2.1 | 0.9 | 15.0 | 0.9 | 2 |
| 20 | 15.4 | 2.2 | 13.5 | 1.6 | 2 |
| 21 | 9.6 | 1.8 | 15.7 | 1.7 | 3 |
| 22 | 4.0 | 1.5 | 14.5 | 6.6 | 3 |
| 23 | 0.9 | 1.8 | 0.6 | 1.9 | 5 |
| 24 | 16.1 | 6.2 | 15.7 | 5.0 | 7 |
| 25 | 15.3 | 2.6 | 12.1 | 1.3 | 7 |
| 26 | 6.4 | 1.8 | 16.1 | 1.4 | 10 |
| 27 | 16.0 | 2.8 | 14.8 | 3.0 | 12 |
| 28 | 4.1 | 1.2 | 15.5 | 2.7 | 17 |
| 29 | 3.4 | 2.6 | 16.1 | 5.7 | 22 |
| 30 | 2.1 | 0.9 | 15.6 | 1.9 | 36 |
| 31 | 12.2 | 1.1 | 13.3 | 0.8 | 40 |

Codebook

id: ID for analysis (sequential from original publication)

ZIKV_s: Optical density (OD) ratio, ZIKV in serum

DENV_s Optical density (OD) ratio, DENV in serum

ZIKV_CSF Optical density (OD) ratio, ZIKV in cerebrospinal fluid

DENV_CSF Optical density (OD) ratio, DENV in cerebrospinal fluid

days Sample collection, days from birth

